# SmartSpacer: Design, Implementation, and In-Vitro Validation of a Multimodal Sensorized Knee Spacer for Continuous Infection Monitoring in Two-Stage Revision Arthroplasty

**DOI:** 10.64898/2026.04.24.720557

**Authors:** Paul Morandell, Christoph Dillitzer, Bach Tran Nguyen, Vincent Lallinger, Igor Lazic, Rainer Burgkart, Oliver Hayden

## Abstract

Periprosthetic joint infection (PJI) is the leading cause of failure in two-stage revision total knee arthroplasty (TKA). The timing of reimplantation currently relies on subjective clinical assessment, as no established method enables continuous, objective, local monitoring of infection dynamics during the spacer interval. We present the SmartSpacer, a sensorized antibiotic-loaded PMMA knee spacer integrating a miniaturized PCB within the tibial component (65 × 45 × 12 mm). The system incorporates digital temperature sensors, a CMOS camera module, a spectrometer, an inertial measurement unit, and a Bluetooth Low Energy (BLE) 5.2 transceiver. Firmware was developed on Zephyr RTOS with aggressive power management. Validation experiments covered power consumption profiling, BLE signal transmission through air, phantom liquid, and ex-vivo porcine knee tissue, temperature accuracy against a calibrated PT100 reference, and motion detection in seven healthy volunteers across three activity protocols. Firmware optimization reduced quiescent current from 700–850 µA to 8 µA, projecting a battery life exceeding 600 days at a clinically relevant sampling rate of one image and one spectrum per hour — more than an order of magnitude beyond the maximum spacer implantation duration. BLE connectivity was maintained reliably up to 6 m through tissue-equivalent phantom liquid and up to 8–9 m in open air. Temperature sensors achieved ±0.16 °C steady-state accuracy with self-heating artefacts below 0.15 °C. Motion detection scaled proportionally with activity intensity, though inter-subject variability in crutch-walking indicated that patient-specific calibration will be required. The SmartSpacer introduces an in vivo wearable - a temporary, implantable knee spacer providing continuous, wireless, multiparametric monitoring within the joint space. It has the potential to transform two-stage revision arthroplasty from empirically timed to data-driven, individualized clinical decision-making.

## 1 Background

### 1.1 Clinical context of total knee arthroplasty and periprosthetic joint infection

Total knee arthroplasty (TKA) is among the most frequently performed orthopedic procedures worldwide, with continuously increasing incidence driven by demographic aging and higher functional demands in both the elderly and younger populations Kurtz et al. (2007). In the United States alone, more than 700000 knee arthroplasties are performed annually, with projections indicating a substantial rise in primary and revision procedures over the coming decades. Carr et al. (2012) Despite the high success rates of primary implantation, revision surgery remains common, particuarly in younger patients, where lifetime revision risks may reach up to 35 % Bayliss et al. (2017).

Periprosthetic joint infection (PJI) represents one of the most severe complications following TKA and is the leading cause of revision surgery. While the overall incidence of PJI is estimated at 1 - 2 %, its clinical and economic burden is disproportionately high Ayoade et al. (2023), Szymski et al. (2024). Septic revisions are significantly more costly than aseptic procedures and are associated with prolonged hospitalization, functional impairment, and increased mortality, with reported one-year mortality rates exceeding 10 % and rising further in elderly populations Zmistowski et al. (2013).

The pathophysiology of PJI is characterized by bacterial colonization of the implant surface and subsequent biofilm formation, which confers resistance to both host immune responses and antibiotic therapy Tande and Patel (2014), Staats et al. (2021). Clically, PJI presents a broad spectrum ranging from acute infections with systemic signs to low-grade infections with subtle, nonspecific symptoms such as persistent pain and reduced joint function Ayoade et al. (2023). These low-grade infections are particularly challenging due to their diagnostic ambiguity and frequent culture-negative results Palan et al. (2019), Parikh and Antony (2016). Figure 1 illustrates the indications and the TKA workflow, highlighting the uncertainty regarding the timing of reimplantation.

**Fig. 1.**
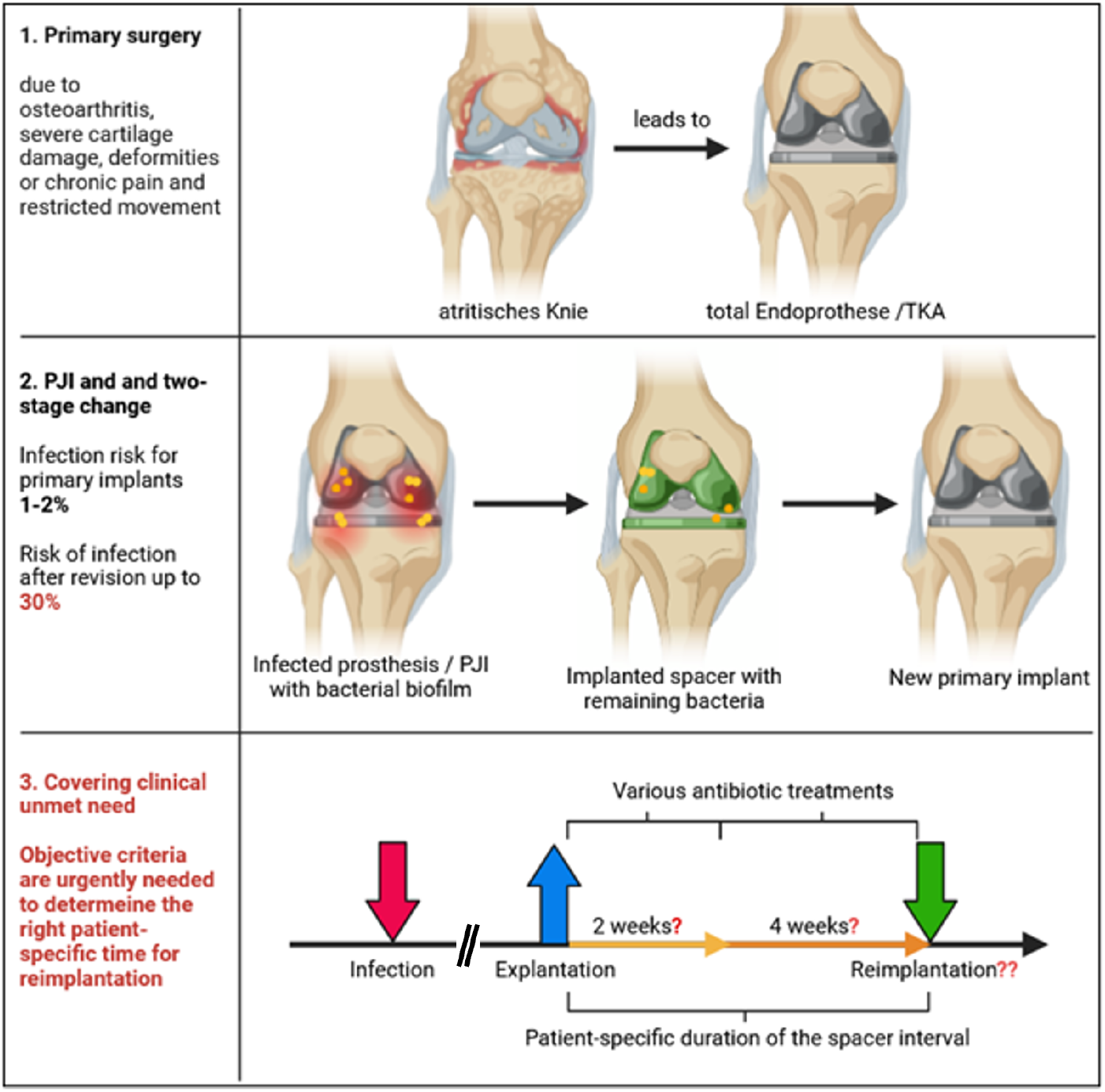
Overview of indications and surgical procedures. 1. Primary surgery (TKA): Primary total knee arthroplasty (TKA) is indicated due to various inflammatory processes as well as degenerative changes of the cartilage and joint structures. 2. Periprosthetic joint infection (PJI) and two-stage revision: The infection rate following primary implantation is approximately 1–2 % ; however, after revision procedures, infection rates increase dramatically and may reach up to 30 % . Following debridement and spacer implantation, residual bacteria may remain within the intraarticular space. The decision regarding the timing of reimplantation is currently the responsibility of the treating surgeons and is largely based on their subjective assessment of clinical criteria. 3. Clinical unmet need / problem statement: This represents a significant clinical unmet need, as both reimplantation timing and patient management during the spacer interval currently lack objective data and reliable local infection markers.

### 1.2 Role and limitations of knee spacers in two-stage revision

The current gold standard for managing chronic or low-grade PJI is the two-stage revision procedure. This involves removal of the infected prosthesis, extensive debridement, and implantation of a temporary spacer, followed by delayed reimplantation of a new prosthesis after presumed eradication of the infection Steinicke et al. (2023). Knee spacers are typically composed of antibiotic-loaded polymethylmethacrylate (PMMA) and serve the purposes of maintaining the joint space, soft tissue tension, and limited mobility as well as local antibiotic delivery. Spacer implantation periods typically range from 2 to 6 weeks but may extend to several months in complex cases. However, antibiotic elution declines significantly after approximately 10 days, limiting long-term antimicrobial effectiveness Batailler et al. (2025), Fink and Tetsworth (2025). A critical unresolved challenge in this treatment paradigm is the lack of reliable, objective, and continuous methods to assess eradication of infection. The current decision-making relies on clinical signs (e.g., swelling, warmth), systemic biomarkers (e.g., CRP), and synovial fluid analysis Al-Jabri et al. (2024), Straub et al. (2024). These approaches are either subjective, nonspecific, or invasive. Consequently, the timing of reimplantation remains uncertain, which can lead to premature reimplantation with persistent infection or prolonged spacer retention with increased morbidity and healthcare costs. To date, no clinically established system enables continuous, in vivo and non-invasive monitoring of local infection dynamics during the spacer phase Puetzler et al. (2023), Li et al. (2021), Straub et al. (2024), Nelson et al. (2023).

### 1.3 Motivation for a sensorized knee spacer

This critical diagnostic gap motivates the development of of a sensorized (“smart”) knee spacer, capable of addressing longitudinal, in situ monitoring of infection-related biomarkers. Conceptually, such a system transforms a passive implant into an active diagnostic platform, which provides continuous data to support personalized clinical decision-making. The feasibility of integrating electronics into implantable medical devices is well established through systems such as cardiac pacemakers, implantable loop recorders, and neurostimulators Joung (2013). These devices demonstrate that long-term implantation of miniaturized electronics, wireless communication, and embedded sensing is technically achievable under strict regulatory and biocompatibility constraints Kim et al. (2024) Jasim et al. (2025). In contrast to permanent implants, the temporary nature of a knee spacer represents a unique opportunity for an in vivo wearable Misir (2025), Allizond et al. (2022):

- reduced requirements for long-term biocompatibility validation
- lower cumulative risk exposure
- greater flexibility in system design and material selection

thus potentially offering a favorable translational pathway for introducing advanced sensing technologies into orthopedic applications Wu et al. (2026). Figure 2 graphically shows the functioning of such a device. Personalized, continuously collected patient data aids the physician in therapeutic decision-making.

**Fig. 2.**
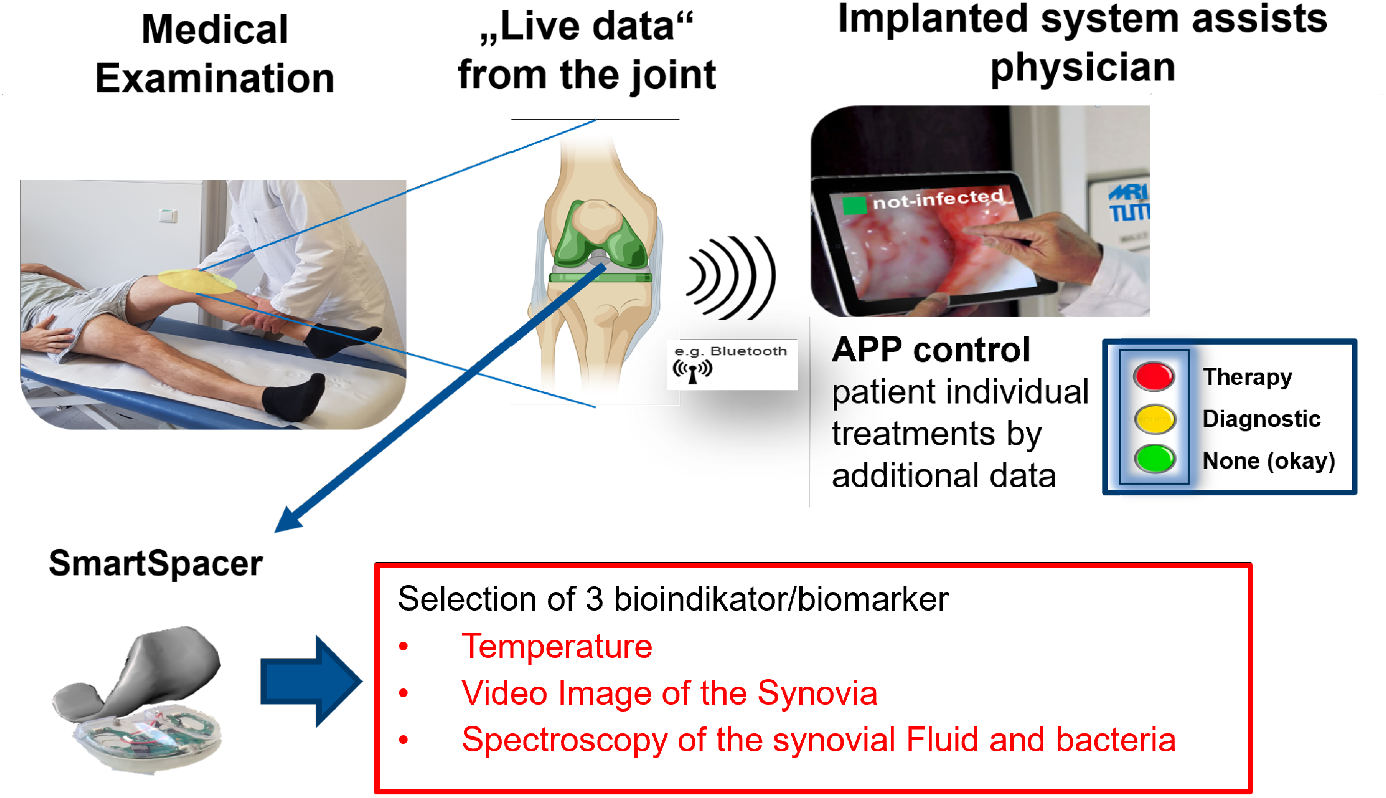
Concept of the digitized knee spacer system. The sensor array is embedded within the tibial component of the spacer. Multiparametric sensor data and endoscopy-free, non-invasive intraarticular images of the synovial fluid are transmitted via low-power Bluetooth to a mobile device to support clinical decision-making.

### 1.4 Design rationale and biomarker-driven component selection for the SmartSpacer

Following the clinical motivation for continuous monitoring during the spacer interval, the next critical step was the translation of clinically relevant infection markers into a technically feasible sensing architecture that can be integrated into a temporary orthopedic implant. As a medical implant, the smart spacer underlies strict constraints. Contrary to a permanent active implant however it was conceived as a temporary, fully encapsulated system, cast intraoperatively and with a consumable character, which allowed for more flexibility in the design. The component selection process was governed by a unique set of boundary conditions that required simultaneous optimization of:

- clinical biomarker relevance
- miniaturization and geometric integrability
- energy efficiency
- cost-effectiveness for potential serial use
- regulatory feasibility and encapsulation safety

The objective was not to maximize sensor complexity, but rather to identify a set of sensing modalities that provide the highest clinical information density per occupied volume, power consumption, and system cost. This resulted in the following main sensing concepts: temperature, optical imaging, and spectral tissue analysis.

#### 1.4.1 Biomarker-driven sensor selection

The sensor selection process was based on a translational engineering principle: only biomarkers that are both clinically meaningful for PJI surveillance and realistically integrable into the limited spacer geometry were considered.

##### Local temperature as an inflammatory biomarker

Local temperature changes represent one of the most robust physiological surrogate markers of inflammation and infection Yishake et al. (2014). As Fajferek et al. (2025) et. al. have shown, it is a relevant biomarker in postoperative infection monitoring. In PJI, local inflammatory responses are associated with immune cell infiltration and activation, while classical inflammatory mechanisms such as vasodilation contribute to the characteristic local increase in temperature and metabolic activity. Furthermore, longitudinal monitoring may enable, alongside absolute temperature elevation, circadian patterns, short-term fluctuations, persistent thermal asymmetries (if temperature is taken on topologically different points), and dynamic response to antibiotic therapy. These temperature-related informations may hold diagnostically relevant data, which makes temperature a valid biomarker for infection monitoring. From an engineering perspective, different types of temperature sensors can be considered. As for this work, it was decided to use digital semiconductor temperature sensors. The many advantages, such as an extremely small footprint, low power consumption, low component cost, high sensitivity, and easy PCB integration, made them a favorable choice. As mentioned above, a key design choice was the placement of multiple temperature sensors to gain a) topologically distributed temperature values close to the tissue, and b) a sensor near the processor to track and compensate for possible self-heating during high load phases.

##### Optical imaging for visual inflammatory assessment

Macroscopic tissue characteristics, including erythema, turbidity, vascular morphology, bleeding, and surface reflectance, may correlate with the underlying inflammatory status and can be captured using an optical, camera-based imaging system Henche and Holder (2012). The integration of a miniaturized camera system was therefore selected to transfer this established visual diagnostic principle into a continuous in vivo monitoring platform. Analogous to the approach employed for temperature sensing, the camera-based modality is designed to enable longitudinal tracking of dynamic changes, particularly those not adequately captured by temperature measurements alone. As outlined above, this primarily includes the progression of tissue erythema, alterations in vascular morphology, as well as changes in surface reflectance patterns and turbidity. From a system design perspective, camera modules offer a highly favorable balance between:

- clinical information richness (correlates with resolution)
- component availability
- miniaturization potential
- cost in large-scale procurement
- power consumption

Compared with more complex imaging modalities, camera integration was considered the most feasible optical approach for a temporary implant.

##### Spectral sensing for microbiological and metabolic characterization

A third biomarker domain of potential interest concerns the microbiological and biochemical activity of the periimplant environment. A major cause of periprosthetic joint infection are bacterial colonization alongside with a biofilm formation and inflammatory metabolite production Costerton et al. (1999). These changes alter the optical scattering and reflectance behavior of surrounding tissue and fluid Al Ghaithi et al. (2024). The longitudinal observation of these properties may again provide diagnostically relevant information. Miniaturized spectrometric sensing was therefore selected to enable indirect assessment of:

- bacterial burden
- biofilm-associated changes
- metabolic activity
- tissue composition changes

In particular, spectral reflectance measurements may provide insight into bacterial concentration and metabolite-associated optical signatures. This modality complements conventional RGB imaging by adding quantitative wavelength-dependent information, thereby increasing diagnostic specificity. Although spectrometers generally impose greater design constraints than temperature sensors, their inclusion was justified by the substantial increase in potential microbiological information content.

##### Why these components were selected

A range of sensing modalities was considered, including fluorescence spectroscopy, ultrasound, and multispectral imaging. Based on the criteria:

1. **Biomarker relevance**: Each selected modality directly maps to clinically established infection indicators. This direct clinical interpretability is essential for later translational acceptance.
2. **Technical feasibility**: The extremely limited implant volume necessitated strict prioritization of components with minimal footprint and simple integration. Further, energy efficiency was of major importance to guarantee the necessary runtime.
3. **Economic scalability**: Because the spacer is a temporary consumable implant, component cost is substantially more relevant than in permanent implant systems. The selected sensor combination provides high diagnostic yield while remaining compatible with economically realistic clinical deployment.

The aim was to achieve an optimal balance between diagnostic value and engineering feasibility, the selection was ultimately limited to these three sensors. Figure 3 provides an overview of potential measurement approaches, their diagnostic value and costs, and the systems ultimately implemented.

**Fig. 3.**
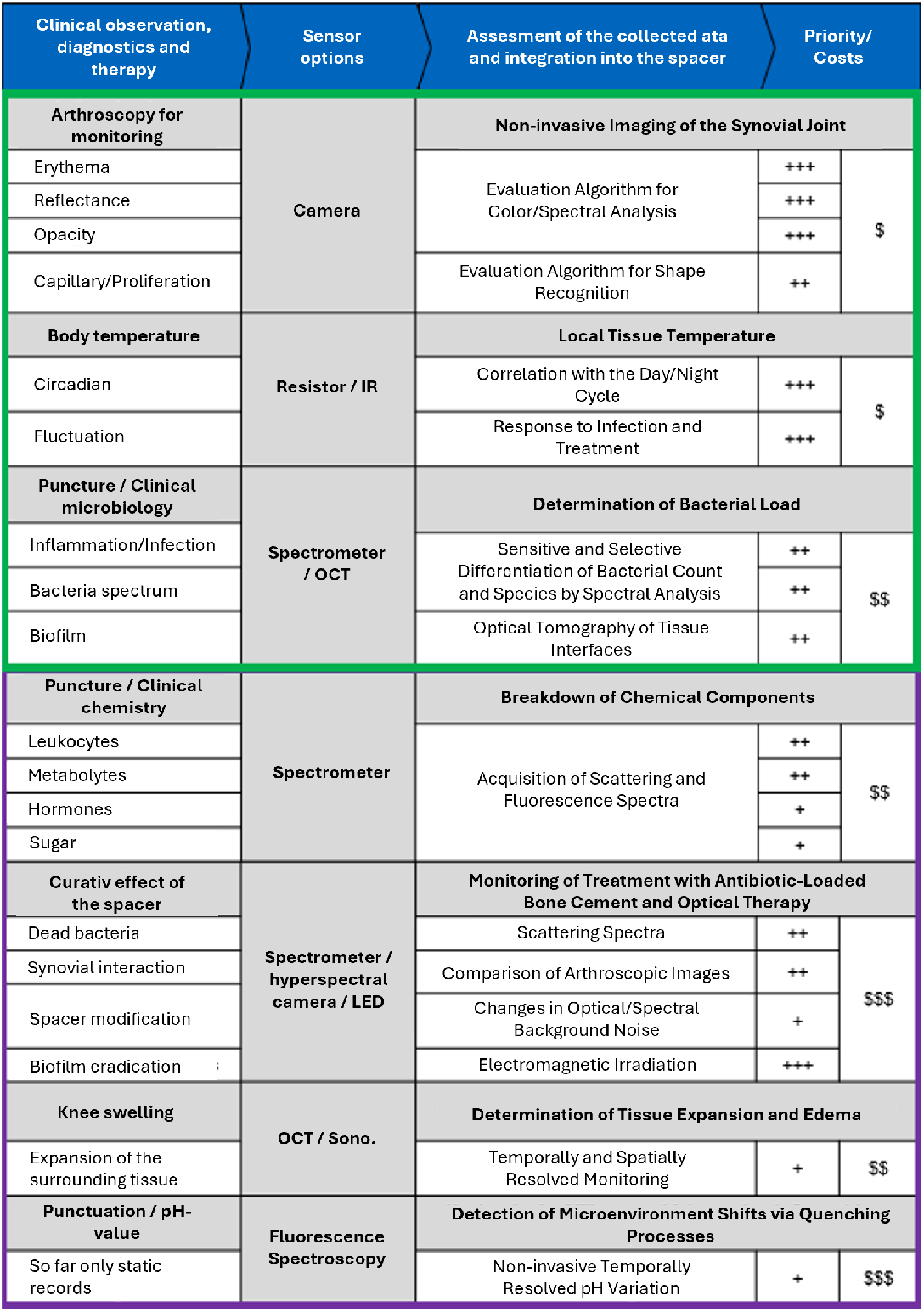
Overview of clinical biomarkers relevant to the diagnosis and treatment of two-stage total joint arthroplasty revision. Specification of corresponding sensor options for the SmartSpacer, including evaluation strategies and integration into the spacer. Prioritization of measurement techniques (low + to high +++) and associated costs (low $ to high $$$) for the overall project. Green framing highlights the most promising detector options.

### 1.5 Geometric constraints as a primary design driver

A central engineering challenge of the SmartSpacer lies in the extremely limited available volume. The tibial spacer component was selected as the primary integration compartment. For the smallest clinically relevant configuration (size S), the available dimensions were:

- width: 65 mm
- length: 45 mm
- height: 12 mm

These dimensions represent a worst-case integration scenario Craig et al. (2022). All components had to fit within this restricted space without compromising structural integrity, mechanical load-bearing behavior, surgical handling, or PMMA encapsulation quality. This geometric limitation strongly influenced all component decisions, particularly regarding battery selection, PCB architecture, and optical pathway design.

#### Optical window as a mandatory geometric feature

Optical sensing introduces an additional geometric challenge: unlike temperature sensors, optical components require a dedicated optical interface. A dedicated PMMA optical window was therefore incorporated into the tibial plateau, serving as the only optical access point to surrounding tissue. Its design was guided by the requirements of high optical clarity, resistance to the PMMA casting process, biocompatibility, and mechanical stability. The dome-shaped window geometry was derived from clinically approved capsule endoscopy systems, providing an established translational pathway Iddan et al. (2000). The angled positioning of approximately 8–10° was as expected to reduce orthogonal reflection artifacts and improve tissue visualization.

#### Miniaturized PCB and energy constraints

The spatial limitations of the spacer directly constrained battery size, which as the largest single component made energy consumption a primary system-level design parameter, requiring all selected components to exhibit low average power demand. A miniaturized multilayer PCB architecture was therefore developed to integrate sensing electronics, signal processing, wireless communication, and power management within a flat cavity of the tibial plateau. Special attention was paid to thermal hotspot avoidance, electromagnetic interference reduction, and reproducible intraoperative positioning. This design strategy ensures that the electronics remain fully encapsulated while maintaining long-term autonomous operation throughout the implantation period, which typically ranges from several weeks to months Batailler et al. (2025), Fink and Tetsworth (2025).

#### Translational significance of the design concept

The final multimodal design represents a deliberate engineering compromise between clinical information content and realistic implant integration. Rather than pursuing maximal sensor complexity, the SmartSpacer focuses on a carefully selected set of biomarkers considered most relevant for monitoring infection dynamics in the clinical context of two-stage revision. This multimodal architecture enables continuous monitoring of inflammation dynamics, visual tissue changes, and microbiological activity, while remaining compatible with the geometric, economic, and regulatory constraints of a temporary implantable orthopedic system.

## 2 Methods

### 2.1 System Architecture and Hardware Design

The SmartSpacer integrates all sensing electronics into the tibial component of a standard articulating PMMA knee spacer (COPAL^®^ mould system, Heraeus Medical GmbH). The printed circuit board (PCB) was designed around the geometric constraints of the smallest clinically relevant tibial plateau (size S: 65 × 45 × 12 mm). A multi-layer PCB architecture accommodates the following main components on a kidney-shaped footprint that conforms to the available tibial cavity:

- **Microcontroller / BLE module:** Nordic Semiconductor nRF5340, dual-core SoC (application core + network core) with integrated Bluetooth Low Energy 5.2 stack. The network core handles wireless communication; the application core executes sensor logic and data management.
- **Camera module:** Osiris M (Optasensor GmbH), a small form-factor CMOS image sensor with a 320 × 320 pixel active matrix (2.4 µm × 2.4 µm pixel pitch, 1/15” optical format, 1.05 mm × 1.05 mm chip size). The sensor operates in free-running rolling-shutter mode with a programmable frame rate of 4–49 fps. Data is transmitted via a two-wire LVDS interface (or optionally single-ended SEIM mode), with power consumption of 12 mW in LVDS mode and 3.2 mW in idle mode. A field of view of 90° is achieved via a custom PMMA dome window angled 8–10° to minimise orthogonal reflection artefacts. An LED ring (4 × white LEDs, 8 mW each, 600 mcd, 130° aperture) provides intraarticular illumination.
- **Spectrometer:** nanoLambda NSP32, an ultra-compact 32 × 32 nano-filter array spectral sensor covering up to 350–1050 nm (device-specific range depending on variant), with a spectral resolution of 10–30 nm and a peak repeatability of 1 nm. The sensor communicates via SPI interface and operates from a 3.3 V single supply, with a typical current consumption of 6 mA during acquisition and 60 µA in power-saving mode. It is individually calibrated per sensor ID. The sensor receives rearward-scattered light through dedicated optical apertures in the camera board.
- **Temperature sensors:** Two Sensirion STS3x-DIS (accuracy ±0.1 °C, I^2^C) and one STS4x (accuracy ±0.2 °C), spatially distributed across the board; one sensor placed adjacent to the MCU to monitor and correct for self-heating.
- **Inertial sensor:** STMicroelectronics ISM330DLC iNEMO™ (3-axis accelerometer + gyroscope; only the accelerometer is used in firmware to conserve energy).
- **Power supply:** 550 mAh LiPo battery (3.6 V), target operating time over six weeks.

To minimise standby current, dedicated P-MOS/N-MOS power switches on the PCB supply the camera and spectrometer only during active acquisition sequences. The complete assembly is cast intraoperatively in epoxy and PMMA bone cement using the standard COPAL^®^ moulding procedure Heraeus Medical GmbH (2024a,b). Figure 4 shows the assembled SmartSpacer PCB with its key components: the nearinfrared spectrometer, the camera module with hemispherical dome for intra-articular imaging, the nRF5340 Bluetooth module, and the ISM330DLC position sensor. Two of the three temperature sensors are indicated on the visible board surface; the third sensor is mounted on the opposite side of the PCB and is therefore not visible in this view. The right side of the figure shows the finite-element simulation of the fully encapsulated spacer, illustrating the top view of the board embedded in bone cement as well as a cross-sectional view revealing the relative positioning of the components within the spacer body.

**Fig. 4.**
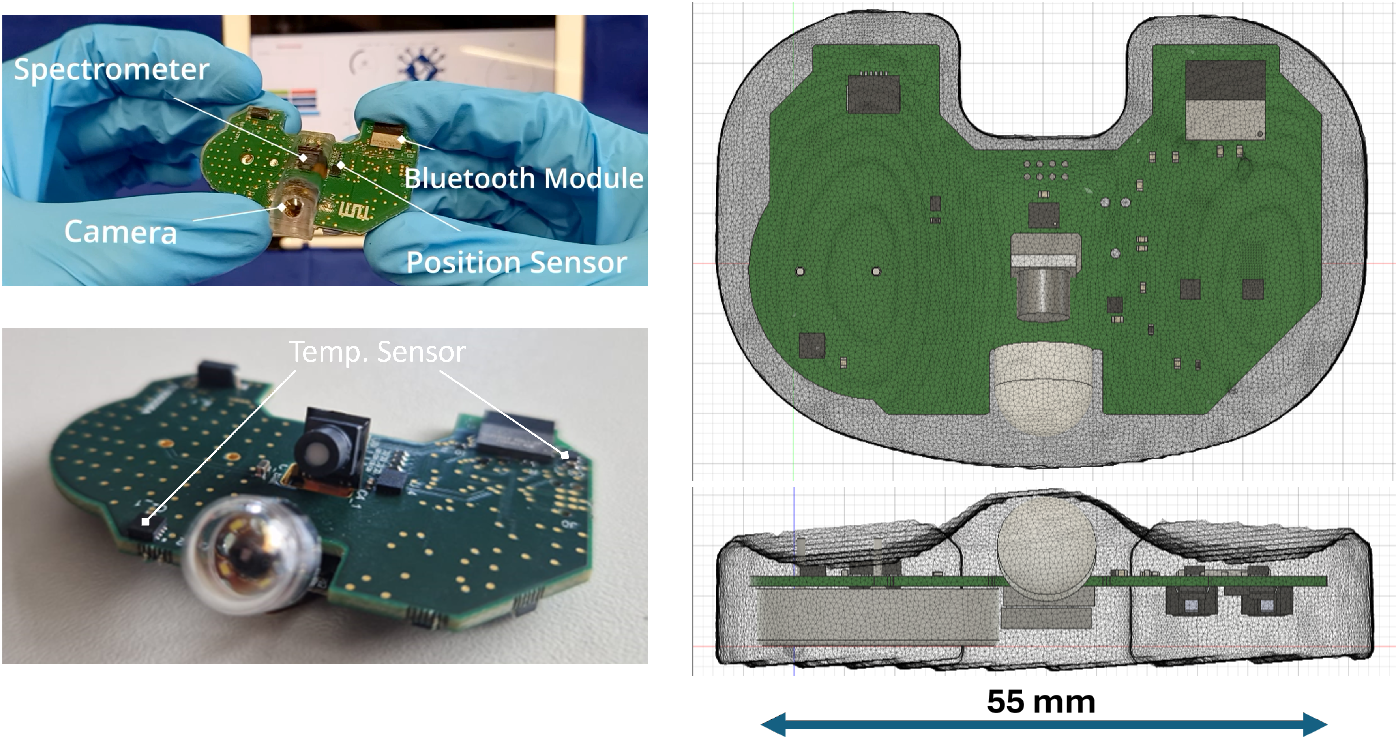
shows the assembled PCB with labeled components (left), the board profile with the hemispherical camera dome (bottom left), and finite-element simulation of the fully encapsulated spacer in bone cement in top and cross-sectional view (right).

### 2.2 Firmware and Power Management

A custom firmware was developed on the Zephyr RTOS using the Nordic Connect SDK (NCS). The primary objective was to reduce idle-state current below 491 µA — the theoretical maximum for 6-week operation with a 550 mAh battery and 10 % capacity degradation:

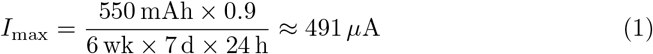

The unoptimised baseline firmware exhibited average currents of 700–850 µA. Profiling with a Nordic Power Profiler Kit II (PPK2, 10 kHz sampling, 200 nA resolution) identified two principal contributors: (i) the Zephyr logging thread and UART console subsystem continuously preventing entry into low-power idle; (ii) large Bluetooth advertising intervals keeping the radio active. Optimisation steps included:

1. Removal of logging, UART/USB-CDC, and serial console on both cores via project configuration flags (CONFIG LOG=n, CONFIG SERIAL=n), allowing the idle thread to execute and the MCU to enter sleep between BLE events.
2. Advertising interval tuning: a 500 ms interval was selected as a compromise between power and reconnection latency.
3. Dynamic connection-parameter management: two BLE connection parameter profiles are defined — a fast profile (7.25 ms connection interval) used during data transfer, and a slow profile with 276 intervals of connection latency (ca. 2 s sleep)
4. 2 Mbit/s PHY: enabled via CONFIG BT USER PHY UPDATE, halving the on-air time per packet.
5. Hardware power-gating: the camera and spectrometer are switched off between acquisition cycles via the P-MOS/N-MOS gate circuit.

After full optimisation, the baseline quiescent current was reduced to 8 µA, with communication events (advertising and connection bursts) producing peaks of up to 5 mA. The time-averaged current during advertising and connected-idle states settled at approximately 30 µA owing to the duty-cycled radio bursts. Image acquisition and BLE transfer draws a mean of 3 mA over ≈ 5.5 s per image at 320 × 320 px. Spectral acquisition and transfer is more demanding due to the higher power draw of the NSP32, the LED illumination ring, and the larger per-measurement data payload, resulting in a mean of 22 mA over ≈ 1.3 s per spectrum. The complete measurement cycle is managed by a state-machine firmware loop: the device advertises at low power, establishes a BLE connection on demand, executes sensor acquisition (temperature, spectrum, image, motion), transfers data to the mobile gateway, and returns to low-power advertising.

### 2.3 Signal Transmission Through Biological Tissue

Because electromagnetic signals are attenuated by water-rich tissue at 2.4 GHz, the wireless range of the implant was characterised under three conditions: open air (reference), phantom liquid, and porcine ex vivo knee. Phantom liquids were prepared from household materials to replicate the dielectric properties of skin (50 % deionised water + 50 % sugar), fat (2.9 % H_2_O + 0.1 % NaCl + 30 % vegetable oil + 67 % flour), and muscle (59.5 % H_2_O + 0.5 % NaCl + 40 % sugar) Kumar and Shanmuganantham (2017) Gabriel et al. (1996). The composition mass was calibrated against CT-based tissue density values of a human knee. A Bluetooth dongle (nRF52840) encapsulated in epoxy resin was submerged in the phantom or surgically implanted in the porcine knee. A mobile receiver (nRF Connect app) was moved in 50 cm steps from 0 to 10 m; RSSI was recorded 10 times per position and averaged. Disconnect frequency and reconnect latency were additionally measured with the production PCB inside phantom liquid using the AIRcable adapter, with 3-minute dwell times per position.

### 2.4 Temperature Sensor Validation

Temperature sensors were validated in a stirred water bath inside a HettCube 200R incubator (37 °C setpoint, ±0.1 °C temporal stability, ±0.2 °C spatial homogeneity). A reference sensor was used simultaneously: a calibrated PT100 RTD (mark D-K-21043-01-00). The SmartSpacer board was fully encapsulated in epoxy and submerged; the three temperature sensors (T1, T2, T3) were recorded every 5 s via BLE. Operating modes (Advertising, Connection, Image Transfer) were systematically cycled to quantify self-heating offsets, and thermal behaviour was additionally modelled in COMSOL Multiphysics 6.1 using the real PCB layout (FR4 + copper multilayer) with measured power dissipation values as boundary conditions.

### 2.5 Motion Detection

Motion detection was based on pitch and roll angles derived from accelerometer data (ISM330DLC; gyroscope disabled for energy reasons). A threshold-based firmware algorithm increments a counter whenever the angular change Δpitch exceeds 10°:

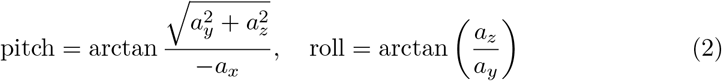

Validation was performed with seven healthy volunteers (n = 7); the PCB was mounted on a 3D-printed knee holder at the distal thigh. Three protocols were evaluated: (1) seated leg extensions at 10, 15, and 30 movements/min (each 60 s × 5 repetitions); (2) walking 10/20/40 steps with 180° turns; (3) the same step protocol with forearm crutches and no weight-bearing on the instrumented leg.

## 3 Results

### 3.1 Power Consumption and Battery Life

After firmware optimisation the quiescent baseline current was reduced to 8 µA, with an average of approximately 30 µA during advertising and connected-idle operation — a reduction of more than 95 % compared to the unoptimised baseline of 700–850 µA. Communication bursts produced transient peaks of up to 5 mA but with low duty cycle. A mean of 3 mA was measured during image acquisition and transfer ( ≈ 5.5 s per image at 320 × 320 px); spectral acquisition and transfer required a mean of 22 mA over ≈ 1.3 s per spectrum, reflecting the combined load of the NSP32, LED ring, and elevated BLE data throughput. System-level energy modelling across realistic usage scenarios is summarised in Table 1. The per-hour current consumption was calculated by integrating the measured current profiles of each operational phase (advertising/idle baseline, image acquisition and transfer, spectral acquisition and transfer) weighted by their respective duty cycles. At a clinically practical sampling frequency of one image and one spectrum per hour, the total consumption of 34.72 µAh/h corresponds to a projected battery life of approximately 600 days — far exceeding the 6-week implantation target by more than an order of magnitude. Even at 30 images and 30 spectra per hour, a runtime exceeding 57 days (≈ 8 weeks) is projected, still meeting the maximum clinical requirement.

**Table 1.** Projected current consumption and battery life (500 mAh) for key usage scenarios.

| Scenario | Consumption ($\mu\text{Ah/h}$) | Battery life (days) |
| --- | --- | --- |
| Advertisement only | 33.4 | 624 |
| Connection, idle | 22.4 | 930 |
| 1 image + 1 spectrum / h | 34.7 | 600 |
| 6 images + 6 spectra / h | 97.2 | 214 |
| 30 images + 30 spectra / h | 362.8 | 57 |
| 60 images + 60 spectra / h | 703.2 | 30 |

### 3.2 Bluetooth Signal Transmission

In open air, BLE signal strength exceeded − 90 dBm across the full 10 m measurement range with zero disconnects recorded at any position (Fig. 5, Fig. 6). Transmission through phantom liquid introduced a mean RSSI attenuation of − 9.1 dBm relative to the in-air reference, while propagation through the porcine ex vivo knee resulted in a lower attenuation of − 5.7 dBm, consistent with the intermediate water content of biological tissue compared to the phantom formulation. Connection stability measurements confirmed reliable connectivity through phantom liquid up to 6 m, with no or negligible disconnects (≤ 1 per 3-minute interval). Beyond 6.5 m, disconnect frequency increased substantially, reaching a mean of 8.5 disconnects per interval at 8.5 m (Fig. 6). In all conditions, automatic reconnection was achieved in *<* 1 s. These results comfortably exceed the connectivity requirements anticipated for clinical use (patient-room bedside distance ≤ 5 m) and the planned porcine preclinical setting (≤ 5.5 m enclosure diameter).

**Fig. 5.**
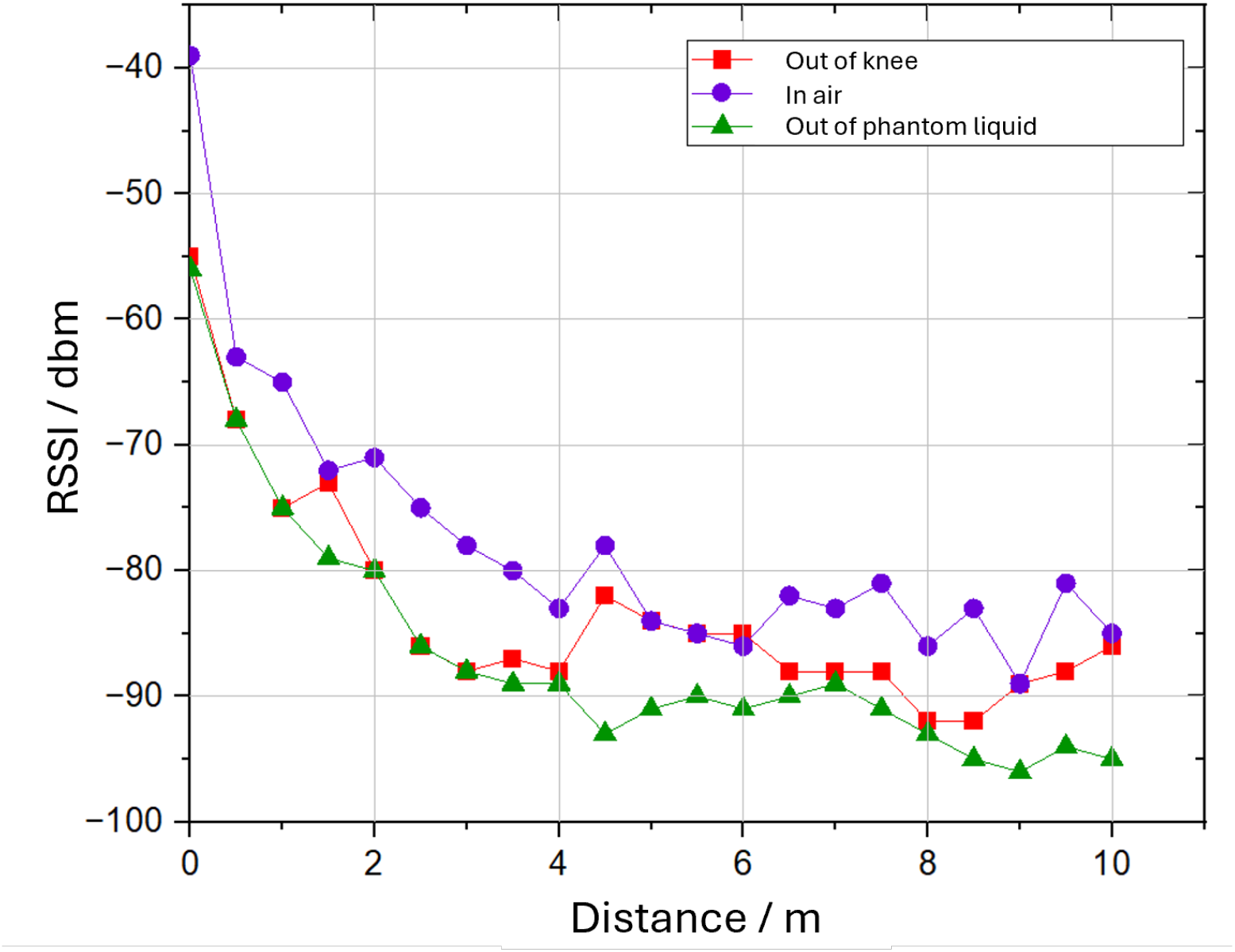
Maximum RSSI as a function of distance for three propagation media. Signal strength decreases monotonically with distance in all conditions, with in-air measurements (red) showing the least attenuation, followed by transmission through the porcine ex vivo knee (blue) and phantom liquid (yellow), reflecting the increasing dielectric losses of each medium.

**Fig. 6.**
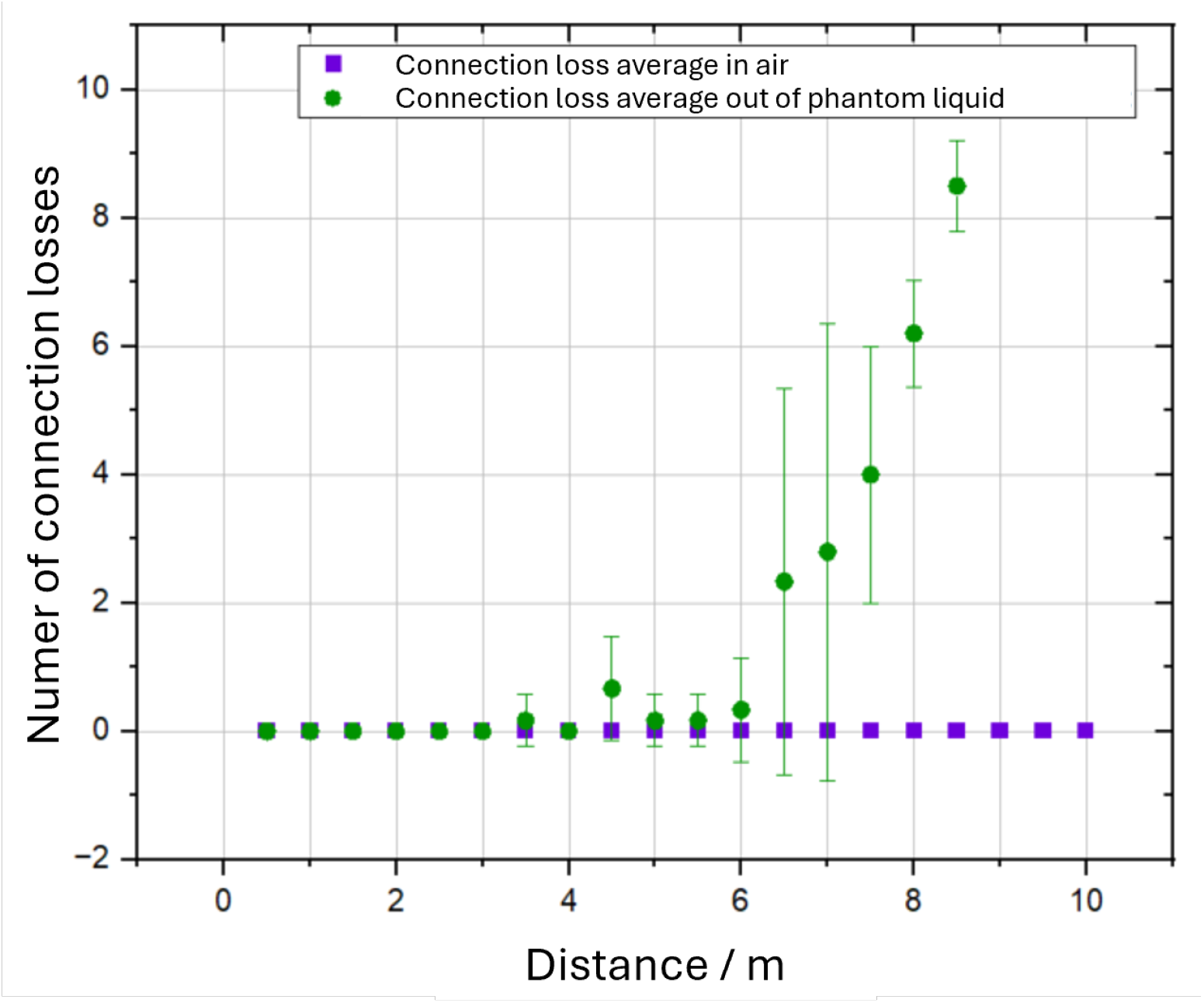
Mean number of disconnects per 3-minute interval as a function of distance, comparing transmission through air (black squares) and through phantom liquid (red circles) using the AIRcable Host XR 4.2 external Bluetooth adapter. No disconnects were recorded through air across the full measurement range. In phantom liquid, connection remained stable up to 6 m, with disconnect frequency increasing substantially beyond 6.5 m. Error bars indicate standard deviation across repeated measurements.

### 3.3 Temperature Measurement

Thermal COMSOL simulations confirmed that self-heating at the three sensor positions remained *<* 0.1 °C in advertising mode and *<* 0.2 °C during peak image transfer. Experimentally measured mode-specific offsets were: idle *<* 0.01 °C; BLE advertising 0.04 – 0.06 °C; BLE connection 0.05 – 0.08 °C; sustained image transfer + 0.12 − 0.15 °C (stabilising within ≈ 3 min of cooldown). These offsets are deterministic and can be corrected in post-processing. In the water-bath validation, the mean temperature of the three sensors converged to within *±* 0.16 °C of the PT100 reference after approximately 3 hours of thermal equilibration. The initial offset of 0.36 °C decreased monotonically as the board reached thermal equilibrium with the medium. Inter-sensor standard deviation was *±* 0.02 °C throughout the experiment, with no measurable influence of cast thickness or proximity to the MCU on steady-state accuracy. The convergence value of 0.16 °C is within the specified sensor accuracy (STS3x *±* 0.1 °C; STS4x *±* 0.2 °C) and is clinically relevant given that PJI-related skin temperature asymmetry decreases from ≈ 5.5 °C on post-operative day 1 to ≈ 1 °C after one month of recovery Yishake et al. (2014).

### 3.4 Motion Tracking

The accelerometer-based angular-change counter correctly scaled with movement intensity in all three protocols. Walking test: angular change counts increased linearly with step count (mean ± SD at 20 steps: 248 *±* 16 across all subjects), confirming proportional detection. Seated extension: the algorithm detected approximately twice the angular-change count when twice as many knee extensions were performed (proportional scaling validated at 10, 15, 30 movements/min). Walking with crutches: individual results were consistent (e.g., proband 4: SD = 13.1 at 20 steps), but inter-subject variance was high (SD = 60.4 at 20 steps) due to gait style differences—experienced crutch users generated up to 80 % more angular changes through limb swing. These findings establish proof-of-concept for motion-context annotation of optical and spectral measurements but indicate that personalised threshold calibration or machine-learning-based gait classification will be required for clinical deployment.

## 4 Discussion

The SmartSpacer concept addresses a well-defined clinical gap: the absence of objective, continuous, local infection monitoring during the inter-stage spacer interval of two-stage TKA revision. The results presented here demonstrate that integrating multimodal sensing electronics into an antibiotic-loaded temporary knee spacer is technically feasible across all critical performance dimensions.

### Energy budget

The optimised system comfortably operates within a 500 mAh battery for 6 weeks at clinically meaningful sampling frequencies. The baseline quiescent current of 8 µA confirms that the combination of aggressive firmware sleep management and hardware power-gating has effectively eliminated residual leakage in standby. The dominant contribution to time-averaged current (≈ 30 µA) stems from duty-cycled radio bursts during advertising and connected-idle states; this is an inherent characteristic of BLE and cannot be further reduced without compromising reconnection latency or reliability. Image acquisition contributes modestly to the energy budget (3 mA × 5.5 s per image), whereas spectral measurements are the more energy-intensive modality (22 mA × 1.3 s per spectrum) due to the simultaneous activation of the NSP32, the LED illumination ring, and the associated BLE transfer of larger data payloads. Despite this, even the most demanding simulated scenario (60 images + 60 spectra per hour) yields a projected runtime of 30 days, which satisfies the minimum clinical spacer interval. Maximising Bluetooth packet size (MTU 251 bytes, Data Length Extension) remains a pending optimisation that will further reduce fractional on-air time per kilobyte transferred.

### Signal transmission

BLE at 2.4 GHz experiences meaningful tissue attenuation (≈ 5–9 dBm through knee-equivalent media), but the system maintains stable connectivity up to 8–9 m, far exceeding the maximum patient-room or bedside distance. Water-based tissue absorbs 2.4 GHz energy effectively Lunkenheimer et al. (2017), making attenuation predictable and reproducible.

### Temperature sensing

The 0.16 °C steady-state precision, achieved with an off-the-shelf digital sensor array, is sufficient to detect the several-kelvin intra-articular temperature elevation characteristic of active PJI and to track its gradual resolution over the treatment course Romano et al. (2011). The small, deterministic self-heating offsets (≤ 0.15 °C during data transfer, resolving within 3 min) are mode-dependent and correctable by firmware. A sensor positioned near the MCU (T3) can serve as a thermal reference to subtract processor self-heating from the other two measurement sensors, a strategy that requires prospective calibration in future animal studies.

### Motion detection

The current threshold-based algorithm provides a binary movement indicator (active vs. resting limb) that may serve as a contextual annotation layer for optical data. The clinical motivation for this annotation rests on the assumption that movement-induced physiological responses — including transient erythema, elevated local perfusion, and increased synovial fluid turbidity — could potentially influence the visual appearance of intraarticular images captured by the camera module, possibly confounding the distinction between mechanically induced hyperaemia and persistent, load-independent inflammatory signs more characteristic of active PJI Pua (2015). Whether such movement-related optical changes are in practice sufficiently pronounced to affect image-based assessment within the enclosed intraarticular space of a spacer-implanted knee remains to be established in future in vivo experiments. Post-operative swelling and synovial fluid dynamics are known to be partly load-dependent, which provides a physiological basis for this hypothesis, but the degree to which these effects manifest as detectable optical signal changes at the sensor level is currently unknown. The motion data therefore serves primarily as a contextual metadata channel at this stage, with its diagnostic utility to be evaluated prospectively. Regarding absolute activity quantification, the high inter-subject variability observed in crutch-walking data indicates that a fixed threshold is insufficient across patients, and that individual baseline calibration or machine-learning-based classification of raw inertial data will likely be required for any future quantitative application.

### 4.1 Limitations

The in-vitro nature of all experiments reported here precludes conclusions about invivo biocompatibility, mechanical durability under physiological loading, or absolute sensor accuracy within the inflammatory synovial milieu. Biomechanical validation against ISO 14879-1, biocompatibility testing (LDH-Glow, Live/Dead assay), and insertion into porcine and ovine ex-vivo specimens have been conducted in parallel (reported elsewhere) and confirm structural integrity and basic biological safety. The animal study phase, currently pending regulatory approval, will provide the first continuous in-vivo dataset.

## 5 Conclusion

We have presented the design, implementation, and in-vitro validation of the SmartSpacer, a sensorised knee spacer prototype integrating temperature measurement, optical imaging, spectroscopy, inertial motion sensing, and Bluetooth Low Energy data transmission. Firmware optimisation reduced idle operating current by more than 60 %, projecting a battery life of at least 20 months at clinically relevant sampling frequencies—exceeding the maximum implantation duration by an order of magnitude. Bluetooth connectivity was maintained reliably up to 8–9 m through tissue-equivalent media. Temperature sensing achieved ±0.16 °C steady-state accuracy with ¡0.15 °C self-heating artefact. Motion detection correctly scaled with activity intensity, though inter-patient calibration remains necessary. Spectral discrimination of bacterial concentrations with the miniaturised implantable spectrometer reached statistically significant performance, with conservative bias towards overestimating rather than underestimating bacterial load—a clinically favourable error mode. Taken together, these results demonstrate that multiparametric, continuous, wireless monitoring of intra-articular infection dynamics from within a temporary knee spacer is technically feasible. The SmartSpacer has the potential to transform two-stage revision arthroplasty from empirically timed to data-driven, patient-individualised clinical decision-making.

## Notes

### Competing Interest Statement

The authors have declared no competing interest.

### Funding

- Bundesministerium für Forschung, Technologie und Raumfahrt: 01EK2107A, 01EK2107B

